# The Physiological Architecture of Groups: Multi-Channel Synchrony Networks Map Individual Perceptions and Collective Behavior

**DOI:** 10.64898/2026.09.18.752699

**Authors:** Alessandro Carollo, Maya Sabag, Nadia Golbez, Ofek Porat, Einav Cohen, Andrea Bizzego, Ilanit Gordon

## Abstract

Humans coordinate in groups, yet much of what we know about coordination comes from dyadic interaction. In groups, however, knowing how much people synchronize may not be enough. Pairwise synchrony can be distributed across teammates: who synchronizes with whom, and how these connections are distributed, may carry distinct social information. Here, we tested whether physiological group synchrony can be understood as a network topology of interpersonal coordination rather than as an aggregate of dyadic alignments. A total of 33 groups (*N* = 161 participants) completed cooperative and competitive joint drumming tasks while cardiac inter-beat intervals (IBI) and electrodermal activity (EDA) were recorded simultaneously. Pairwise physiological synchrony was estimated using lagged cross-correlation and used to construct synchrony-based networks. IBI-based networks showed stronger and broader cardiac connectivity during cooperation, with cardiac coupling also increasing during competitive drumming relative to baseline. EDA-based networks also showed task-sensitive changes in node-level connectivity and captured social and behavioral dimensions of group interaction, linking trust and cohesion to node-level connectivity, and collective behavioral synchrony to lower clustering and higher local efficiency. Together, these findings show that the social meaning of physiological group synchrony lies not only in how much individuals align, but in how physiological alignment is distributed across people, relationships, and biological systems. By preserving this configuration, synchrony-based network analysis captures information about group experience that is lost when synchrony is reduced to an average dyadic score, providing a route toward understanding group synchrony as a flexible, selective, and distributed property of social coordination.

## Introduction

Groups organize social behavior across animal species. Fish gather in schools, birds fly in flocks, ungulates move in herds, and primates form complex social communities. Across these diverse systems, collective living provides important benefits, including protection, information sharing, and collaboration (Krause & Ruxton, 2002; Ward & Webster, 2016).

A key feature of group life is that social organization is not random. Individuals often associate preferentially with some partners, avoid others, and occupy distinct positions within the social network. One mechanism underlying this organization is homophily, whereby individuals who share similar characteristics are more likely to affiliate with one another than with individuals who differ from them (Fu, Nowak, Christakis, & Fowler, 2012; Lewis, Gonzalez, & Kaufman, 2012; McPherson, Smith-Lovin, & Cook, 2001). For instance, across primates, similarities in personality traits may increase interpersonal attraction and facilitate the formation of closer social ties (Massen & Koski, 2014; Nelson, Thorne, & Shapiro, 2011). Importantly, homophily may extend beyond observable behavioral characteristics. Similarity in biological activity across individuals may also contribute to perceived social closeness, suggesting that social bonds are shaped not only by what people do, but also by how their neurophysiological systems respond to each other and the environment (Parkinson, Kleinbaum, & Wheatley, 2018; Shen et al., 2025).

Beyond pre-existing similarity, social interactions can also generate similarity over time. In this sense, synchrony can be understood as a temporal analogue of homophily: whereas homophily captures similarity in relatively stable individual characteristics or response profiles, synchrony describes how similarity emerges and fluctuates dynamically during interaction. In fact, decades of research in interpersonal psychology and social neuroscience have shown that interacting individuals tend to align their behaviors and biological activity during social exchange. This phenomenon, often described as bio-behavioral synchrony, has been observed across species and across multiple modalities, including behavior, physiology, neural activity, and hormonal fluctuations (Feldman, 2012, 2017; Kingsbury et al., 2019; Rose, Styr, Schmid, Elie, & Yartsev, 2021). Importantly, synchrony across these modalities appears to be associated with interpersonal closeness and with the quality of the interaction (Bizzego et al., 2019; Carollo, Bizzego, et al., 2025; De Felice et al., 2025; Djalovski, Dumas, Kinreich, & Feldman, 2021; Endevelt-Shapira & Feldman, 2023; Gordon & Bartsch, 2026; Nguyen et al., 2020). In particular, at the physiological level, interpersonal synchrony has been linked to individual, social, and performance-related outcomes, including affective states, interpersonal processes, and task performance (Gordon et al., 2020).

Most research on synchrony has so far focused on interacting dyads. Although this approach has been highly informative, it overlooks the fact that individuals are embedded within broader social networks, where multiple relationships can shape both individual behavior and the dynamics of other relationships (Gordon, 2025). For this reason, research on interpersonal relationships is increasingly moving toward the study of larger social groups (Esposito & Carollo, 2026). For instance, studies on three-person groups have shown that physiological synchrony is associated with team performance and may contribute to group cohesion (Elkins et al., 2009; Tomashin, Gordon, & Wallot, 2022). Similarly, hyperscanning studies have examined interpersonal neural synchrony in larger social settings, showing that group-level neural alignment is related to classroom engagement, classroom social dynamics, and team performance in adult groups (Dikker et al., 2017; Reinero, Dikker, & Van Bavel, 2021). These findings are consistent with a flexible view of bio-behavioral synchrony (Gordon, 2025; Gordon, Tomashin, & Mayo, 2025), according to which synchrony is not uniformly high across all individuals or contexts, but rather dynamically adjusted to the demands of the social situation. This perspective is especially relevant for group settings, where multiple individuals, roles, and relational demands coexist within the same interaction. In such contexts, effective coordination may require not only alignment, but also differentiation, as group members may need to synchronize with some partners or signals while remaining distinct from others.

Building on this view, our recent theoretical work has proposed that group synchrony should be understood in terms of relational multiplicity: a structured organization of multiple synchrony relations that can vary across individuals, modalities, and interaction contexts (Carollo & Gordon, 2026). This framework shifts the focus from whether a group is synchronized to how synchrony is organized across individuals, relationships, modalities, and relational contexts. From this perspective, group synchrony is not a uniform state shared equally by all members, but a structured and context-dependent configuration of interpersonal connections. The shift from dyads to groups therefore requires a conceptual and methodological extension of existing approaches. In dyadic research, synchrony is typically quantified as the degree of temporal alignment between two interacting individuals. In this context, being “in sync” or “out of sync” necessarily refers to the specific dyad. In groups, however, synchrony is not confined to a single dyadic relationship: it may vary across pairs, subgroups, modalities, and task demands. Thus, group synchrony cannot be fully captured by a single dyadic measure, nor by aggregating all dyadic measures within the group. Instead, it requires methods that can describe how bio-behavioral alignment is organized across the broader social structure. These considerations motivate a network-based approach, because networks can represent not only the magnitude of synchrony, but also how synchrony is arranged across the group.

Social network analysis has become a powerful framework for quantifying social organization (Borgatti, Mehra, Brass, & Labianca, 2009; Carollo & Gordon, 2026). By representing individuals as nodes and their relationships as edges, this approach makes it possible to characterize both direct interpersonal ties and the wider architecture of social systems. In the context of group synchrony, network analysis provides a way to represent synchrony as a coordination topology: a structured pattern of connections that can vary in strength, distribution, and organization across the group (Carollo & Gordon, 2026). This framework preserves pairwise synchrony information and avoids aggregation while embedding each dyadic connection within the broader group structure, making it possible to identify central or peripheral individuals, selectively connected members, local subgroups, and more globally integrated patterns of coordination. This is important because group synchrony may not be adequately represented by averaging pairwise synchrony values across all dyads. Instead, its social and behavioral significance may depend on how synchrony is selectively distributed across individuals and organized within the broader group topology.

Social networks have typically been constructed from self-reported closeness, observed affiliation, or behavioral interaction scores. Far less is known about whether networks can be derived directly from bio-behavioral synchrony itself, and whether such synchrony-based networks capture meaningful aspects of social organization and collective behavior in groups. This gap is particularly important for physiology, because physiological synchrony may reveal forms of interpersonal coordination that are not fully accessible through self-report or overt behavior alone. Physiological signals can capture continuous, embodied changes in arousal and regulation as group members respond to one another during interaction. Deriving networks from these signals therefore makes it possible to ask whether the physiological organization of a group carries social and behavioral information beyond explicitly reported relationships or observable patterns of affiliation.

In the present study, we provide an empirical test of this physiological synchrony-based network framework in groups of five individuals engaged in a shared drumming task designed to manipulate cooperation and competition. We derived physiological synchrony networks from pairwise coupling in inter-beat intervals (IBI) and electrodermal activity (EDA), and used social network analysis to characterize both individual-level and group-level organization of physiological synchrony. At the individual level, we first examined whether node-level network metrics were associated with participants’ subjective perceptions of group functioning. At the group level, we then tested whether the topology of physiological synchrony networks was associated with collective behavioral synchrony. Finally, we examined whether synchrony-based network metrics varied across task contexts involving cooperation and competition. To our knowledge, this is among the first empirical studies to operationalize physiological group synchrony as a synchrony-based network. This approach shifts the question from the magnitude of synchrony between two individuals to the organization of synchrony relations within the group. Physiological group synchrony can therefore be represented as a coordination topology: a structured pattern of interpersonal connections that captures who is connected to whom, how strongly, and how these connections are organized across the group. By preserving this relational structure, synchrony-based networks make it possible to test whether physiological group synchrony carries information about individual experience and collective behavior that would be obscured by averaging pairwise synchrony across dyads.

## Methods

### Experimental design

The study used a repeated-measures group interaction design to examine how different forms of social coordination and competition influenced physiological synchrony in small groups. Groups of four or five participants completed a baseline period followed by four joint drumming conditions: cooperative drumming, harmonious drumming, in-group competition, and out-group competition. Cardiac activity and EDA were recorded simultaneously from all group members, and behavioral coordination during drumming was captured from MIDI data. After each condition, participants also completed self-report measures assessing their perception of the group.

Pairwise physiological synchrony was used to construct IBI- and EDA-based social networks, from which individual- and group-level network metrics were derived. These metrics were then used to test their associations with participants’ perceptions of the group, their relationship with behavioral synchrony during the drumming task, and their sensitivity to experimental condition.

The study was conducted in accordance with the ethical principles of the Declaration of Helsinki and was approved by the Human Subjects Institutional Review Board at Bar-Ilan University (protocol no. 221224397).

### Participants

A total of 161 participants, organized into 33 groups, took part in the study. Groups consisted of four or five participants. Participants had a mean age of 26.52 years (*SD* = 7.16, range = 19-55 years). Of the 161 participants, 105 identified as female, 53 as male, one identified with another gender, and two did not report their gender.

Of the initially recruited groups, one group (*n* = 5 participants) was excluded from both the IBI and EDA analyses because of recording difficulties that prevented reliable reconstruction of the physiological signals. Two additional groups (*n* = 10 participants) were excluded from the EDA analyses because of errors during data storage, and one group was excluded from the behavioral synchrony analyses because of errors in performance detection (*n* = 5 participants). All participants provided informed consent prior to participation.

### Experimental tasks

The experimental procedure was based on a joint drumming task. This task was selected because it is nonverbal, thereby facilitating the acquisition of high-quality neurophysiological data while minimizing speech-related artifacts. In addition, joint drumming has been used in previous work as a goal-oriented paradigm to examine the emergence of social engagement and coordination among non-drummer participants (D’Ausilio, Novembre, Fadiga, & Keller, 2015; Fairhurst, Janata, & Keller, 2013; Gordon et al., 2020; Keller, Novembre, & Hove, 2014). After receiving a detailed explanation of the study procedure and providing informed consent, participants were fitted with electroencephalography (EEG) caps, as well as electrodes placed on the torso and palms to record electrocardiographic and electrodermal activity.

The procedure followed a fixed order 1. Participants first completed a 5-minute baseline recording, during which they were instructed to sit quietly with their eyes open and to avoid moving, talking, or making eye contact. Following the baseline, groups completed four active drumming conditions designed to manipulate the degree and target of collaboration and competition. After each experimental condition, participants completed brief self-report questionnaires assessing their subjective experience of the preceding task in relation to the group. These measures are described in detail in the “Individual perception of the group” section. Each active condition lasted 4 minutes and was administered in the following order: cooperative drumming, harmonious drumming, in-group competition, and out-group competition. Participants then completed two control clapping conditions, each lasting 2 min and 20 s.

In the cooperative drumming condition, participants were instructed to work together to find as many joint rhythms as possible. Once the group identified a shared rhythm, they were asked to drum it for a short period and then, when they felt that the rhythm had been exhausted, to move on and search for a new rhythm. This condition was designed to encourage participants to repeatedly synchronize and desynchronize behaviorally while pursuing a shared group goal.

In the harmonious drumming condition, participants were instructed to be as creative and harmonious as possible in their drumming. They were encouraged to generate as many harmonious drum patterns as possible, without necessarily playing the same rhythm at the same time. Thus, this condition emphasized complementary coordination and musical coherence rather than explicit behavioral synchrony.

In the in-group competition condition, participants were told that each group member should try to lead as many drumming patterns as possible and that individual performance would be evaluated based on the number of rhythms each person led. Although the instructions did not explicitly frame the task as a competition among group members, the manipulation was designed to encourage competition for leadership within the group.

In the out-group competition condition, participants were first told that, in a previous drumming round, their group had produced a certain number of rhythms, a performance comparable to that of most groups, but that some groups had produced a greater number of rhythms and patterns. They were then given another opportunity to improve their performance and try to outperform other groups. This condition was designed to induce competition at the group level, shifting the competitive focus from individual leadership within the group to collective performance against other groups. Finally, during the two control clapping conditions, participants were instructed to keep their eyes open and to refrain from making eye contact with one another. They listened to a pre-recorded audio track of an audience clapping, in which the clapping pattern alternated between periods of synchrony and desynchrony. In the present study, however, analyses focused only on the baseline and the four active drumming conditions.

### Acquisition and pre-processing of physiological data

Throughout the experiment, participants’ cardiac activity was recorded using electrocardiography (ECG), together with their EDA (see Figure 2A). The following subsections describe the acquisition and pre-processing procedures for both physiological signals.

**Fig. 1:**
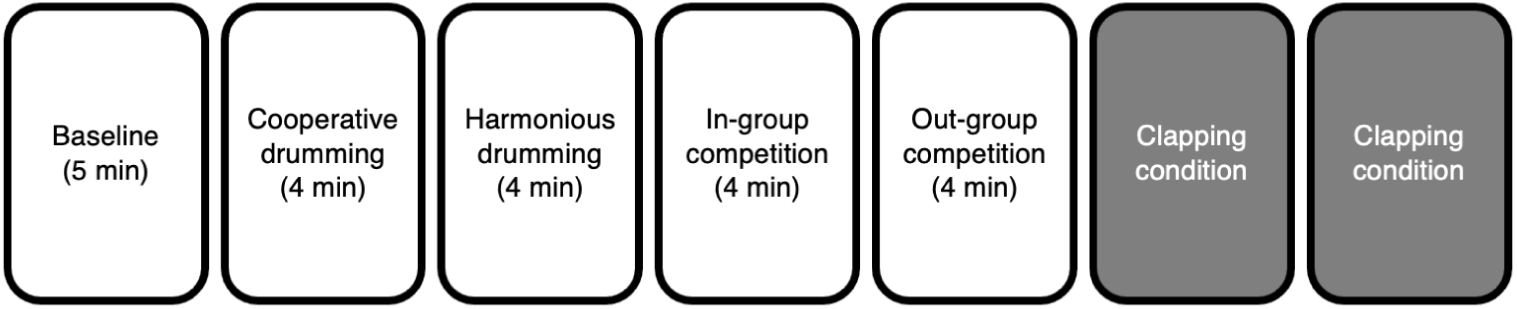
Experimental procedure. Participants first completed a 5-minute baseline recording, followed by four active drumming conditions administered in a fixed order: cooperative drumming, harmonious drumming, in-group competition, and out-group competition. Each active drumming condition lasted 4 minutes and was followed by self-report questionnaires assessing participants’ perceptions of the group. Two clapping control conditions (in gray) were completed at the end of the procedure but were not included in the present analyses.

**Fig. 2:**
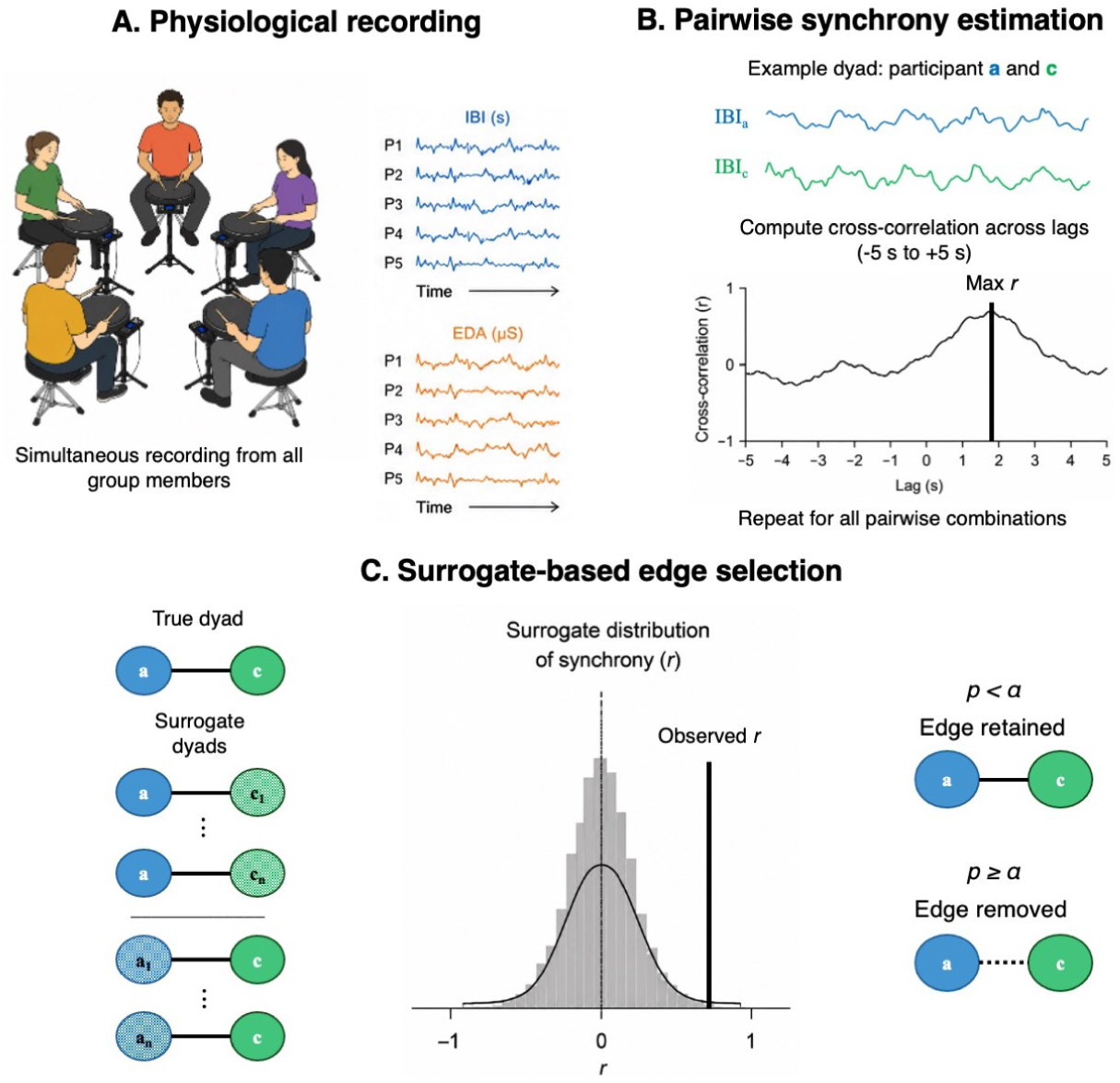
Construction and characterization of physiological synchrony networks. **(A)** Inter-beat interval (IBI) and electrodermal activity (EDA) were recorded simultaneously from all members of each group during the joint drumming task. **(B)** Pairwise interpersonal synchrony was estimated for each dyad using lagged cross-correlation across a *±*5 s window, retaining the maximal cross-correlation value as the synchrony index. **(C)** For each observed dyad, synchrony was compared against a surrogate distribution obtained by pairing each participant with members of other groups, thereby disrupting the original co-presence relationship. Edges were retained only when observed synchrony exceeded surrogate-based expectations after Bonferroni correction.

#### Electrocardiogram

Cardiac activity was recorded continuously from all group members using a modified lead-II electrocardiogram configuration (Sherwood et al., 1990). Electrodes were connected to MindWare Mobile Recorders (MindWare Technology, Gahanna, OH), which synchronously and wirelessly transmitted the ECG signals to a control-room laptop at a sampling rate of 500 Hz. ECG data were processed using MindWare Heart Rate Variability Analysis Software (v3.1.4). Signals were amplified with a gain of 1,000 and filtered using a Hamming windowing function. Trained researchers visually inspected the recordings and manually corrected artifacts and ectopic beats following established guidelines (Berntson et al., 1997). IBI were then extracted from the continuous ECG signal for each participant. The resulting IBI time series were interpolated at 4 Hz to obtain evenly sampled signals, time-aligned across participants, and trimmed to the common overlapping segment across participants. Finally, the time series were standardized and pre-whitened before computing interpersonal synchrony. Pre-whitening was applied to reduce temporal autocorrelation and minimize the possibility that synchrony estimates reflected shared autocorrelated structure in the physiological signals rather than genuine interpersonal temporal coupling.

#### Electrodermal activity

EDA was recorded from each participant at a sampling rate of 500 Hz using two Ag/AgCl electrodes attached to the palm of the nondominant hand. EDA data were processed using MindWare Technologies’ EDA application software (Version 3.1.5). Trained researchers visually inspected the recordings and manually corrected artifacts. The signal was smoothed using a rolling filter with 500 data points per block. Analyses focused on the overall skin conductance signal, including both tonic skin conductance level and phasic skin conductance responses. Given that the task was not event-related and that we had no specific hypotheses concerning distinct tonic and phasic mechanisms, the signal was not detrended or decomposed into separate components. This choice is consistent with previous work on EDA synchrony during continuous social interaction, where the primary aim is to examine overall covariation in sympathetic arousal rather than responses to discrete events (Gordon, Gilboa-Schechtman, Gilboa, Cohen, & Kivity, 2023). The skin conductance signal was downsampled to 2 Hz by averaging values within 500-ms bins. Values were expressed in microsiemens, and a minimum threshold of 0.05 *µ*S was applied to exclude floor-level noise, following the default MindWare setting and prior research (Gordon, Horesh, et al., 2021). As for the IBI time series, EDA time series were time-aligned across participants, trimmed to the common overlapping segment within each dyad or group, standardized, and pre-whitened before computing interpersonal synchrony.

### Interpersonal physiological synchrony

Cross-correlation was used to quantify the presence of shared temporal patterns in both the IBI and EDA time series across group members. Cross-correlation estimates the degree to which two physiological signals co-vary over time (Kettunen & Ravaja, 2000; Kettunen, Ravaja, & Keltikangas-Järvinen, 2000; Mauss, Levenson, McCarter, Wilhelm, & Gross, 2005), while allowing for imperfect temporal alignment between them. Specifically, the inclusion of a lag parameter makes it possible to account for anticipatory or delayed physiological responses in one dyad member relative to the other. In the present study, cross-correlation was computed across multiple lags ranging from -5 s to +5 s, in 0.25-s increments for IBI and 0.5-s increments for EDA. The maximum value across this lag window was then retained as the index of physiological synchrony between the two time series. We refer to this measure as the maximal cross-correlation within a time shift of *±* 5 s. For each group, we computed cross-correlations for all possible pairwise combinations of group members (see Figure 2B).

To obtain a surrogate distribution of synchrony values, we generated surrogate dyads for each pairwise connection of interest. Specifically, for each observed dyad, each of the two members was separately matched with another random participant from each group, thereby creating a sample of surrogate synchrony estimates that preserved the individual participants’ physiological time series while disrupting the original pairwise connection. This surrogate distribution was then used to inform edge selection in the physiological synchrony networks, as described in the following section (see Figure 2C).

### Social network generation

Each group was initially represented as a weighted network based on the pairwise physiological synchrony values computed among all group members (Newman, Barabási, & Watts, 2011). In these networks, nodes represented individual participants, while edges represented the maximal cross-correlation values between each pair of participants.

To identify meaningful synchrony-based connections, each observed edge was compared against the surrogate distribution generated for the corresponding dyad. Specifically, for each true dyad, surrogate dyads were created by matching each of the two dyad members with a randomly selected participant from each of the other groups. This procedure generated a reference distribution of synchrony values that reflected the expected level of physiological synchrony in the absence of the co-presence relationship (Golland, Arzouan, & Levit-Binnun, 2015; Golland, Keissar, & Levit-Binnun, 2014).

For each observed edge, a statistical comparison was then performed between the true pairwise synchrony value and its corresponding surrogate distribution using a one-sample *t*-test. The resulting *p*-values were corrected for multiple comparisons at the group level using the Bonferroni procedure to guarantee a conservative approach (*α* = 0.050*/*10 = 0.005; see Figure 2C). Only edges that survived this correction were retained in the final physiological synchrony networks, separately for IBI and EDA.

### Social network metrics

Social network metrics were extracted at two levels of analysis: the individual level and the group level (see Figure 3).

**Fig. 3:**
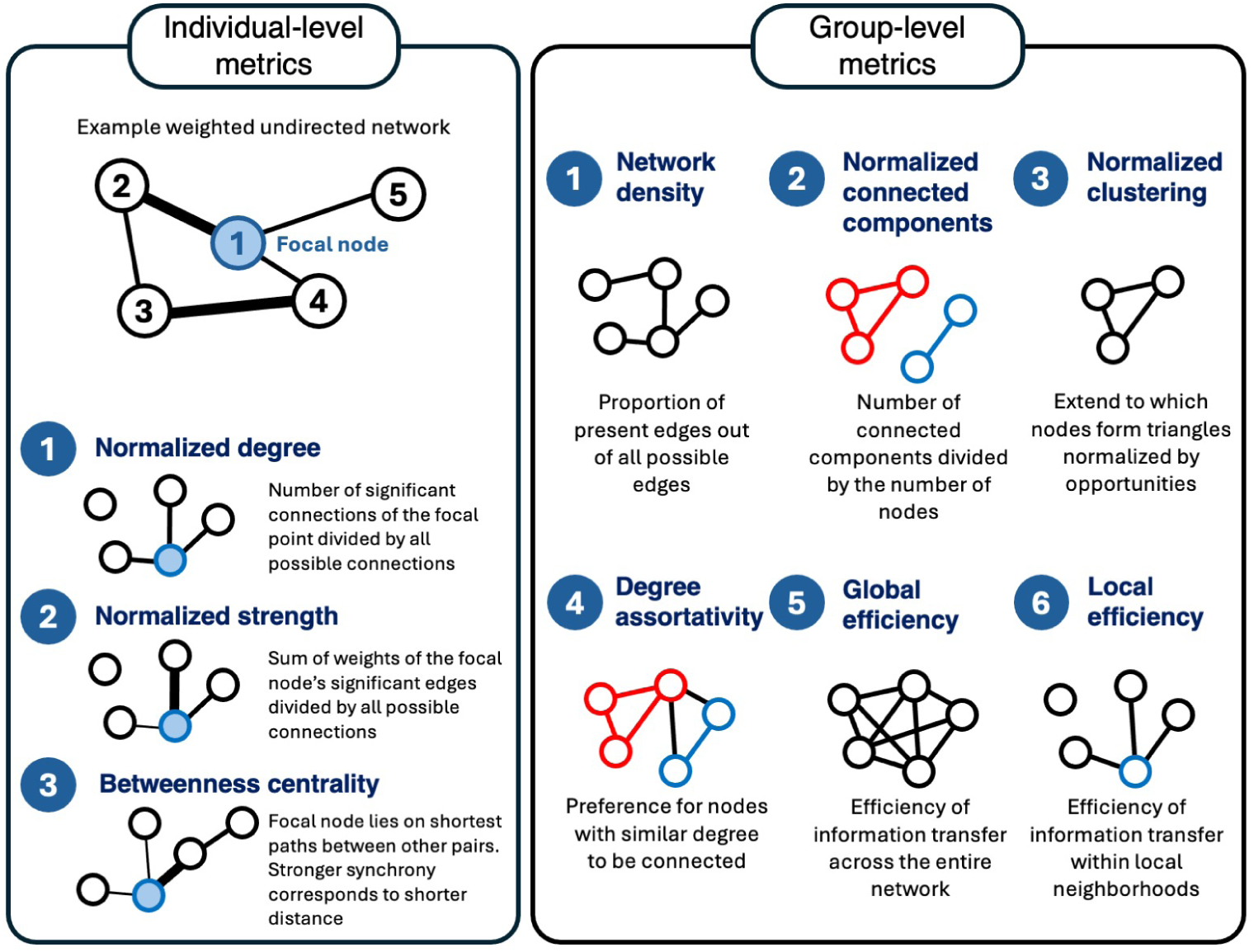
Individual and group-level metrics extracted from the generated social networks.

Metrics describing the individual level included normalized degree, normalized strength, and betweenness centrality. Normalized degree was computed as the number of significant synchrony-based connections involving a given individual, divided by the maximum number of possible connections for that individual. This metric captures the proportion of group members with whom each participant showed significant physiological synchrony. Node strength was computed as the sum of the weights of all significant edges connected to a given individual. Normalized strength was then obtained by dividing node strength by the maximum number of possible connections for that individual, thereby capturing the overall magnitude of physiological synchrony linking each participant to the rest of the group. Finally, betweenness centrality was used to assess the extent to which each participant occupied a bridging position within the physiological synchrony network. Since edge weights represented synchrony strength, they were transformed into inverse weights (1*/w*) before computing betweenness centrality, so that stronger synchrony corresponded to shorter network distances.

At the group level, we extracted network metrics describing the overall structure of physiological synchrony within each group. These included the number of nodes, the number of edges, network density, normalized connected components, normalized clustering, degree assortativity, global efficiency, and local efficiency. The number of nodes corresponded to the participants included in each group network, whereas the number of edges indicated the number of significant synchrony-based connections retained after edge selection. Network density quantified the proportion of observed edges relative to the total number of possible edges, providing an index of how broadly physiological synchrony was distributed across the group. Normalized connected components were computed as the number of connected components minus one, divided by the maximum possible number of components minus one. This metric ranged from 0 to 1, with 0 indicating a fully connected network and 1 indicating a fully disconnected network. Normalized clustering was computed by dividing the weighted average clustering coefficient by network density, thereby indexing the tendency of participants to form locally interconnected synchrony-based triads while accounting for the overall number of edges in the network. Degree assortativity quantified the tendency of nodes with similar degree values to be connected to one another. Finally, global and local efficiency were used to characterize the integration of the synchrony network: global efficiency captured how efficiently coordination could be distributed across the whole network, whereas local efficiency captured the efficiency of coordination within local neighborhoods of connected participants.

### Individual perception of the group

Participants’ subjective perceptions of the group were assessed after each experimental condition using a set of self-report questionnaires. These measures were selected to capture multiple dimensions of participants’ experience of group functioning, including perceived cohesion, collective efficacy, creative performance, trust, group effectiveness, group satisfaction, and leadership. Together, they allowed us to test whether individual differences in the subjective experience of the group were associated with participants’ position and connectivity within the physiological synchrony networks.

Unless otherwise specified, item responses were summed to obtain one score for each construct, participant, and condition, with higher scores indicating higher levels of the corresponding construct. Internal consistency was assessed using Cronbach’s *α* for each multi-item scale.

Perceived group cohesion was assessed using five items rated on a 1-to-6 Likert scale ranging from 1 (not at all) to 6 (extremely). The items captured participants’ perceptions of teamwork, cooperation, mutual reliance, mutual support, and willingness to participate again with the same group. Specifically, participants indicated the extent to which they agreed that they worked together as a team, cooperated with one another, could count on one another, supported one another, and would be happy to participate in another group experiment with the same group. Internal consistency was excellent (Cronbach’s *α* = 0.90).

Collective efficacy was assessed using seven items rated on a 7-point Likert scale ranging from 1 (very little) to 7 (very much). The items captured participants’ confidence in the group’s ability, expertise, and capacity to perform difficult tasks, as well as perceived task interdependence among group members. Specifically, participants indicated the extent to which they had confidence in the group’s ability to perform the task, perceived the group’s level of expertise as high, would feel good about performing a difficult task with the group, needed information and consultation from other group members, depended on them to complete their work, needed to work closely with them, and perceived that other group members needed information from them. Internal consistency was good (Cronbach’s *α* = 0.83).

Perceived creative performance was assessed using three items rated on a 7-point Likert scale ranging from 1 (extremely little) to 7 (extremely much). The items captured participants’ evaluation of the group’s work in terms of innovation, adaptability, practicality, originality, and creativity. Specifically, participants rated how innovative and practical the group’s work was, how adaptive and practical it was, and how creative it was. Internal consistency was good (Cronbach’s *α* = 0.89).

Trust in the group was assessed using seven items rated on a 7-point Likert scale ranging from 1 (very little) to 7 (very much). The measure combined one item assessing group identification (“I identify with my group”) with six items assessing trust and rapport among group members. These items captured participants’ perceived ability to speak freely with other group members about task-related problems, expectations of caring and helpful responses from the group, perceptions of professionalism and dedication, perceived trustworthiness of the group, and trust in group members. Internal consistency was excellent (Cronbach’s *α* = 0.95).

Group effectiveness was assessed using a single item rated on a 7-point Likert scale: “In my opinion, the overall effectiveness of my group was?”. The scale ranged from 1 (not effective) to 7 (very effective), with higher scores indicating greater perceived group effectiveness.

Group satisfaction was assessed using a single item rated on a 7-point Likert scale: “How satisfied are you with your group’s work?”. The scale ranged from 1 (not at all satisfied) to 7 (very satisfied), with higher scores indicating greater satisfaction with the group’s work.

Leadership was assessed after each drumming condition using a peer-nomination item. Participants were asked: “In the present drumming phase, did you recognize a group member that was a clear leader? If so, who?” They could nominate one group member or indicate that there was no leader. For each participant, a leadership score was computed by summing the number of times they were nominated as the leader and dividing this value by the number of members in the group. Higher scores therefore indicated that a participant was more frequently perceived as a leader by the group. When applicable, items were adapted from previous work on group cohesion, collective efficacy, and group processes (Craig & Kelly, 1999; Gordon et al., 2020; Henry, Arrow, & Carini, 1999; Hodges & Carron, 1992; McAllister, 1995; Podsakoff & MacKenzie, 1994). Other items were developed for the present task to capture condition-specific subjective experience.

### Behavioral group synchrony

Drumming performance was analyzed from MIDI recordings using Ableton Max for Live and Max/MSP. Following the approach used in previous work (Gordon et al., 2020), the temporal interval between individual drum hits was extracted and compared across group members to identify temporally coordinated responses. Hits produced by different participants within a fixed temporal window of 30 ms were considered synchronous.

Because groups in the present study consisted of five participants, synchrony was quantified not only for instances in which all group members played simultaneously, but also for partial synchrony patterns involving different numbers of participants. Specifically, synchronous events involving two, three, four, or five group members were identified separately. To account for the number of individuals contributing to each synchronous event, we computed a weighted synchrony sum by multiplying the number of synchronous patterns by the number of participants involved in each pattern. Thus, events involving two participants were weighted by 2, those involving three participants by 3, those involving four participants by 4, and those involving all five participants by 5.

A group-level behavioral synchrony score was then computed by relating this weighted synchrony sum to the total number of drum hits produced during the task. Specifically, the final behavioral measure was calculated as the weighted sum of synchronous events divided by the total number of drum hits. This measure was used as the group-level index of behavioral group synchrony in the subsequent analyses.

### Statistical analysis

Statistical analyses were conducted at both the network and individual levels using linear mixed-effects models (Bates, Mächler, Bolker, & Walker, 2015). All analyses described below were conducted separately for IBI-based and EDA-based physiological synchrony networks. Linear mixed-effects models were chosen because they provide a flexible framework for analyzing hierarchically structured or nested data, while simultaneously accounting for both fixed and random effects. These models are also well suited to datasets with unequal numbers of observations across participants or networks, including cases in which some data points are missing or vary across measurement levels (e.g., Carollo et al., 2026; Carollo, Stella, Lim, Bizzego, & Esposito, 2025).

First, we examined whether individual-level network metrics were associated with participants’ perceptions of the group, separately for IBI- and EDA-based node-level networks. These self-reported measures captured participants’ subjective experience of group functioning, including perceived group cohesion, collective efficacy, creative performance, trust in the group, group effectiveness, group satisfaction, and leadership. Self-report predictors were standardized prior to analysis. The node-level models were extended by including the group-perception measures as fixed-effect predictors, while experimental condition was included as a random effect. Random intercepts for group and participant nested within group were included to account for the hierarchical structure of the data. The in-group competition drumming condition was excluded from these analyses because it introduced competition among members of the same group, thereby altering the nature of the within-group relations targeted by the self-report measures. Including this condition could therefore have reduced the comparability of constructs such as perceived cohesion and trust across conditions. For these individual-level analyses, Benjamini–Hochberg correction was applied separately within each physiological modality and for each group-perception measure across the three node-level metrics: normalized degree, normalized strength, and betweenness centrality. Thus, for each perceptual measure, the three corresponding tests across node-level outcomes constituted a separate family of comparisons.

As a subsequent step, we examined whether the organization of physiological synchrony networks was associated with group-level behavioral synchrony during the drumming task. Separate linear mixed-effects models were fitted for IBI- and EDA-based networks, with behavioral synchrony as the dependent variable. Network density, normalized number of connected components, normalized clustering, degree assortativity, global efficiency, and local efficiency were entered simultaneously as fixed-effect predictors. Experimental condition and group were included as random intercepts to account for condition-related variability and repeated observations within the same group. Because this analysis involved a single multivariable model for each physiological modality, no additional correction for multiple comparisons was applied to this set of predictors.

As a third step, we investigated the effect of experimental condition on synchrony-based network metrics. Separate models were fitted for each group-level network outcome, with experimental condition included as a fixed effect and group included as a random intercept to account for the non-independence of repeated observations within the same group. For individual-level node metrics, participant-level variability was additionally accounted for by including participant as a random intercept nested within group. Thus, individual-level models included experimental condition as a fixed effect, with random intercepts for group and participant nested within group. To account for multiple testing, Benjamini–Hochberg corrections were applied separately within each physiological modality and level of analysis: one correction was applied across the group-level network metrics and a separate correction across the individual-level node metrics. Significant main effects of experimental condition were followed up with pairwise comparisons of estimated marginal means. For each significant outcome, Benjamini–Hochberg correction was applied across the corresponding set of pairwise comparisons.

## Results

### Social network metrics and individual perception of the group

In the first analysis we examined whether individual differences in participants’ perceptions of the group were associated with their position and connectivity within the physiological synchrony networks. Specifically, we tested whether self-reported measures of group functioning, including perceived cohesion, collective efficacy, creative performance, trust, group effectiveness, group satisfaction, and leadership, were related to individual-level network metrics.

For IBI-based synchrony networks, none of the individual group-perception measures were significantly associated with normalized node degree, normalized node strength, or betweenness centrality after Benjamini-Hochberg correction for multiple comparisons (*q* s *>* 0.050).

For EDA-based synchrony networks, several associations with individual perceptions of the group survived Benjamini-Hochberg correction. Perceived trust was positively associated with normalized node degree (*β* = 0.053, *SE* = 0.022, *t* (421.12) = 2.44, *q* = 0.023; Figure 4A). The normalized degree model explained 1.8% of the variance through fixed effects alone (marginal *R*^2^ = 0.018) and 19.9% when both fixed and random effects were considered (conditional *R*^2^ = 0.199). For normalized node strength, perceived trust was positively associated with node strength (*β* = 0.012, *SE* = 0.003, *t* (416.97) = 3.47, *q* = 0.002; Figure 4B), whereas perceived cohesion was negatively associated with node strength (*β* = -0.008, *SE* = 0.003, *t* (412.42) = -2.67, *q* = 0.023; Figure 4C). The normalized strength model explained 2.7% of the variance through fixed effects alone (marginal *R*^2^ = 0.027) and 25.9% when both fixed and random effects were considered (conditional *R*^2^ = 0.259). No significant associations were observed for betweenness centrality, and no other associations with normalized node degree or normalized node strength survived correction for multiple comparisons (*q* s *>* 0.050).

**Fig. 4:**
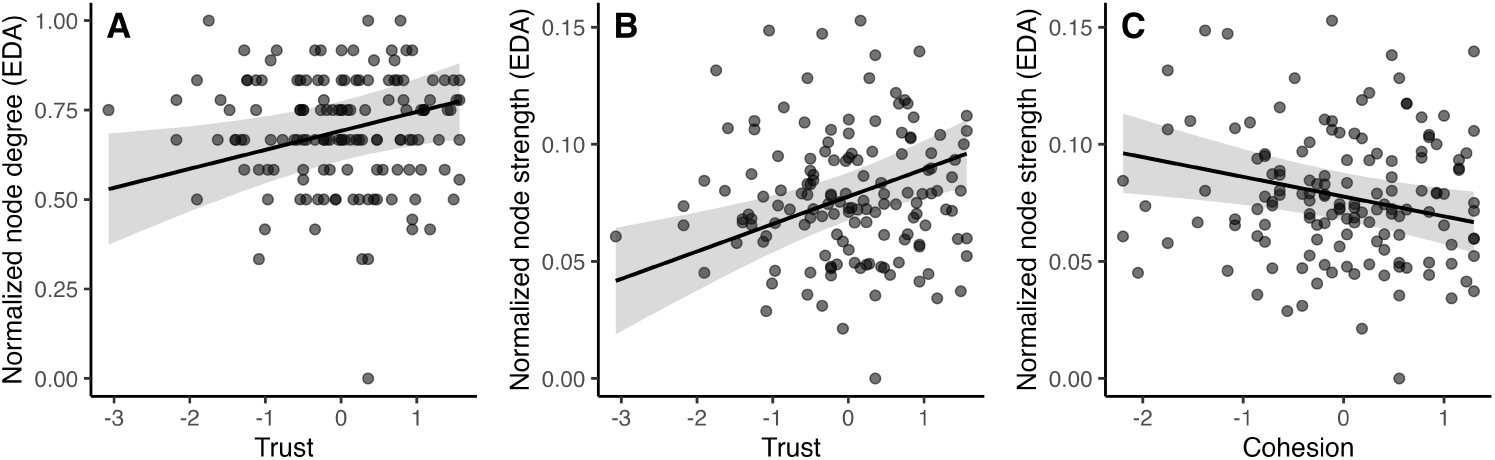
Associations between individual perceptions of group functioning and electrodermal activity (EDA)-based node-level synchrony metrics. Points represent participant-level averages across the three active drumming conditions included in these analyses. Lines and shaded areas show model-estimated fixed-effect associations with 95% confidence intervals, with all other predictors held at their mean values. **(A)** Perceived trust was positively associated with normalized node degree. **(B)** Perceived trust was positively associated with normalized node strength. **(C)** Perceived cohesion was negatively associated with normalized node strength.

### Social network metrics and group behavior

We next examined whether group-level physiological synchrony network properties were associated with behavioral synchrony during the drumming task (Table 1).

**Table 1:** Associations between physiological synchrony network metrics and group behavioral synchrony. IBI = inter-beat interval; EDA = electrodermal activity. (* *p <* 0.050).

| Signal | Network metric | $\beta$ | $SE$ | $df$ | $t$ | $p$ |
| --- | --- | --- | --- | --- | --- | --- |
| IBI | Density | -0.027 | 0.125 | 56.86 | -0.22 | > 0.050 |
| IBI | Normalized connected components | 0.055 | 0.134 | 55.59 | 0.41 | > 0.050 |
| IBI | Normalized clustering | 0.018 | 0.036 | 71.80 | 0.49 | > 0.050 |
| IBI | Degree assortativity | -0.008 | 0.017 | 65.13 | -0.46 | > 0.050 |
| IBI | Global efficiency | 0.037 | 0.185 | 53.39 | 0.20 | > 0.050 |
| IBI | Local efficiency | -0.003 | 0.064 | 70.44 | -0.06 | > 0.050 |
| EDA | Density | 0.046 | 0.121 | 62.51 | 0.38 | > 0.050 |
| EDA | Normalized connected components | -0.176 | 0.122 | 65.20 | -1.45 | > 0.050 |
| EDA | Normalized clustering | -0.039 | 0.016 | 60.95 | -2.41 | **0.019 \*** |
| EDA | Degree assortativity | 0.002 | 0.011 | 64.65 | 0.18 | > 0.050 |
| EDA | Global efficiency | -0.173 | 0.192 | 64.75 | -0.90 | > 0.050 |
| EDA | Local efficiency | 0.068 | 0.027 | 58.88 | 2.53 | **0.014 \*** |

For IBI-based synchrony networks, none of the network-level metrics were significantly associated with group behavioral synchrony (*p*s *>* 0.050).

For EDA-based synchrony networks, normalized clustering was negatively associated with group behavioral synchrony (*β* = -0.039, *SE* = 0.016, *t* (60.95) = -2.41, *p* = 0.019; Figure 5A), whereas local efficiency showed a positive association (*β* = 0.068, *SE* = 0.027, *t* (58.88) = 2.53, *p* = 0.014; Figure 5B). The model explained 14.43% of the variance through fixed effects alone (marginal *R*^2^ = 0.144) and 59.30% when both fixed and random effects were considered (conditional *R*^2^ = 0.593). Thus, greater behavioral synchrony during drumming was associated with lower clustering but higher local efficiency in EDA-based physiological synchrony networks. No significant associations were observed for density, normalized connected components, degree assortativity, or global efficiency (*p*s *>* 0.050).

**Fig. 5:**
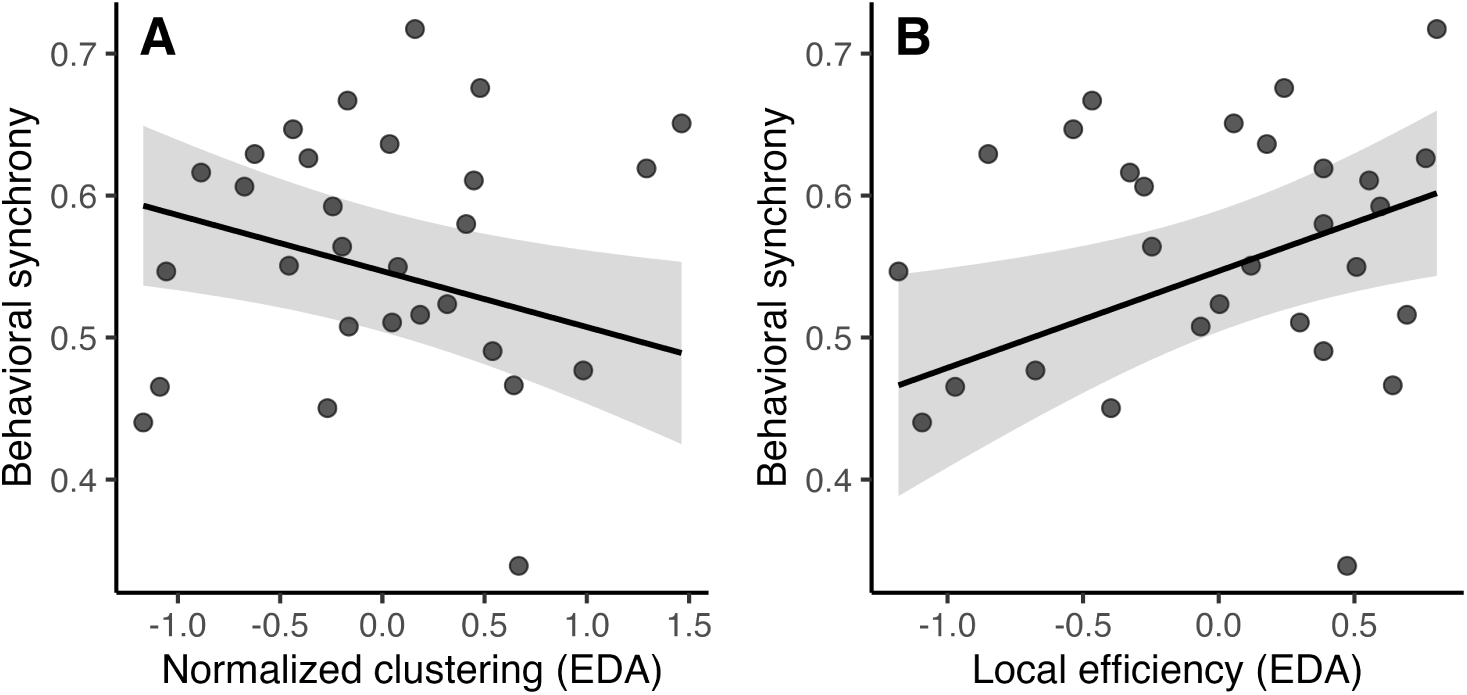
Associations between electrodermal activity (EDA)-based group-level synchrony network metrics and behavioral synchrony. Points represent group-level averages across the four active drumming conditions. Lines and shaded areas show model-estimated fixed-effect associations with 95% confidence intervals from the linear mixed-effects model, with the remaining network metrics held at their mean values. Higher behavioral synchrony was associated with lower normalized clustering **(A)** and higher local efficiency **(B)**.

### Social network metrics across experimental conditions

Finally, we investigated the effect of experimental condition on individual- and group-level network metrics using a series of linear mixed-effects models, separately for IBI- and EDA-based networks. Full model results are reported in Table 2.

**Table 2:** Effects of experimental condition on individual- and group-level physiological synchrony network metrics. *q*-values refer to Benjamini-Hochberg-corrected *p*-values. IBI = inter-beat interval; EDA = electrodermal activity. (** *q <* 0.010, *** *q <* 0.001).

| Signal | Level | Metric | Effect | NumDF | DenDF | $F$ | $q$ |
| --- | --- | --- | --- | --- | --- | --- | --- |
| IBI | Node | Normalized degree | Condition | 4 | 729.66 | 4.74 | **0.001 \*\*** |
| IBI | Node | Normalized strength | Condition | 4 | 729.97 | 9.80 | **<0.001 \*\*\*** |
| IBI | Node | Betweenness centrality | Condition | 4 | 608.00 | 1.37 | $> 0.050$ |
| IBI | Network | Density | Condition | 4 | 155.00 | 1.64 | $> 0.050$ |
| IBI | Network | Normalized connected components | Condition | 4 | 155.00 | 0.24 | $> 0.050$ |
| IBI | Network | Normalized clustering | Condition | 4 | 153.00 | 0.79 | $> 0.050$ |
| IBI | Network | Degree assortativity | Condition | 4 | 105.42 | 0.14 | $> 0.050$ |
| IBI | Network | Global efficiency | Condition | 4 | 155.00 | 0.70 | $> 0.050$ |
| IBI | Network | Local efficiency | Condition | 4 | 155.00 | 1.03 | $> 0.050$ |
| EDA | Node | Normalized degree | Condition | 4 | 572.00 | 6.00 | **<0.001 \*\*\*** |
| EDA | Node | Normalized strength | Condition | 4 | 686.23 | 2.45 | $> 0.050$ |
| EDA | Node | Betweenness centrality | Condition | 4 | 572.01 | 0.39 | $> 0.050$ |
| EDA | Network | Density | Condition | 4 | 116.00 | 2.80 | $> 0.050$ |
| EDA | Network | Normalized connected components | Condition | 4 | 116.00 | 1.01 | $> 0.050$ |
| EDA | Network | Normalized clustering | Condition | 4 | 144.00 | 2.52 | $> 0.050$ |
| EDA | Network | Degree assortativity | Condition | 4 | 98.07 | 0.55 | $> 0.050$ |
| EDA | Network | Global efficiency | Condition | 4 | 116.00 | 2.32 | $> 0.050$ |
| EDA | Network | Local efficiency | Condition | 4 | 145.00 | 2.05 | $> 0.050$ |

For IBI-based synchrony networks, experimental condition had significant main effects on normalized node degree (*F* (4, 729.66) = 4.74, *q* = 0.001, *η*^2^ = 0.03; Figure 6A) and normalized node strength (*F* (4, 729.97) = 9.80, *q <* 0.001, *η*^2^ = 0.05; Figure 6B), whereas no significant effect was observed for betweenness centrality (*F* (4, 608.00) = 1.37, *q >* 0.050). Benjamini–Hochberg-corrected *post-hoc* comparisons indicated that normalized node degree was higher in the cooperative drumming condition than in baseline (*q* = 0.006, *d* = 0.36), harmonious drumming (*q* = 0.001, *d* = 0.45), in-group competition (*q* = 0.014, *d* = 0.32), and out-group competition (*q* = 0.003, *d* = 0.40). No other pairwise differences in normalized node degree were significant (*q* s *>* 0.050). For normalized node strength, cooperative drumming showed higher values than baseline (*q <* 0.001, *d* = 0.70), harmonious drumming (*q <* 0.001, *d* = 0.47), in-group competition (*q* = 0.005, *d* = 0.35), and out-group competition (*q* = 0.005, *d* = 0.36). In addition, normalized node strength was higher during in-group competition (*q* = 0.005, *d* = 0.35) and out-group competition (*q* = 0.005, *d* = 0.34) than during baseline. No other pairwise differences in normalized node strength were significant (*q* s *>* 0.050).

**Fig. 6:**
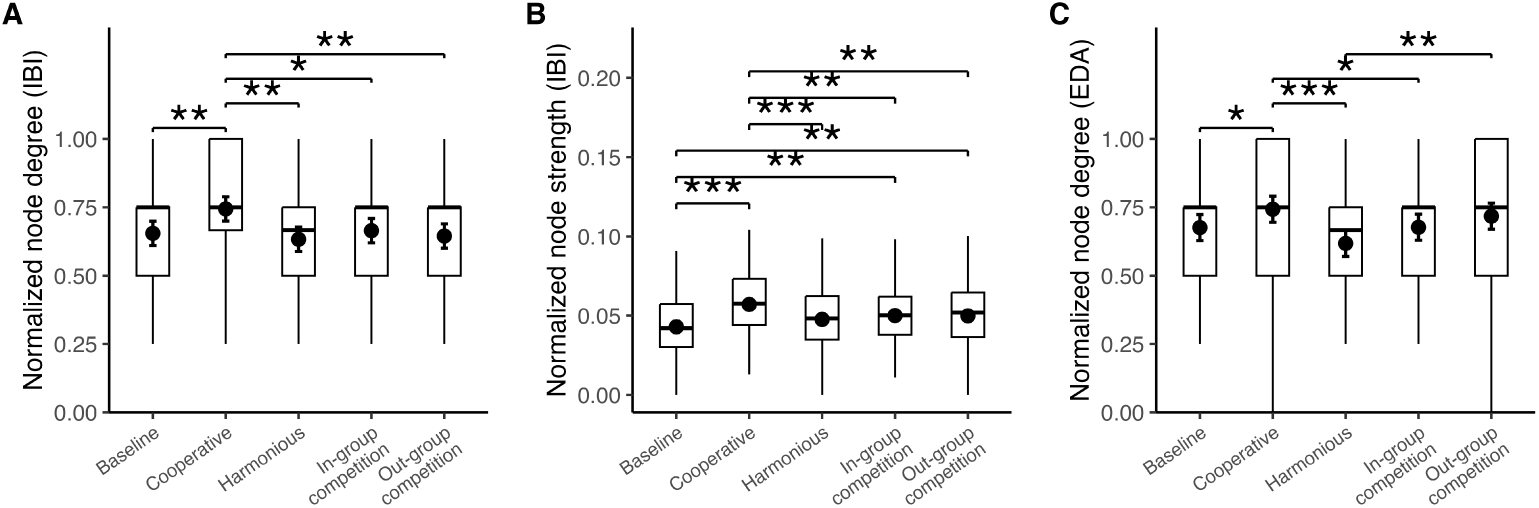
Experimental condition effects on node-level physiological synchrony network metrics. Boxplots show participant-level distributions across experimental conditions. Black points and error bars indicate estimated marginal means and 95% confidence intervals from the linear mixed-effects models. **(A)** In IBI-based networks, normalized node degree was higher in the cooperative drumming condition as compared to the others. **(B)** In IBI-based networks, normalized node strength was highest during cooperative drumming. Strength was higher in the cooperative, in-group competition, and out-group competition conditions than in baseline. **(C)** In EDA-based networks, normalized node degree varied across active drumming conditions, with lower values in the harmonious condition relative to the cooperative and out-group competition conditions, and higher values in the cooperative condition relative to baseline and in-group competition. Asterisks indicate significant Benjamini–Hochberg-corrected *post-hoc* comparisons: \**q <* 0.050, \*\**q <* 0.010, \*\*\**q <* 0.001.

At the network level, experimental condition did not significantly affect network density, the normalized number of connected components, normalized clustering, degree assortativity, global efficiency, or local efficiency (*q* s *>* 0.050). Thus, although IBI-based synchrony showed condition-related changes at the node level, these effects did not translate into large-scale changes in network topology. In particular, node strength increased during active drumming, especially in the cooperative condition, whereas node degree increased mainly in the cooperative condition relative to the other conditions. This pattern suggests that the manipulation modulated how strongly and how broadly individuals were embedded in the IBI-based synchrony network, while the overall architecture of the network remained stable across conditions.

For EDA-based synchrony networks, experimental condition had a significant main effect on normalized node degree (*F* (4, 572) = 6.00, *q <* 0.001, *η*^2^ = 0.04; Figure 6C), whereas no significant effects were found for the other node-level metrics examined (*q* s *>* 0.050). Benjamini-Hochberg-corrected *post-hoc* comparisons indicated that normalized node degree was lowest in the harmonious drumming condition. Specifically, normalized degree was significantly lower in the harmonious drumming condition than in the cooperative drumming condition (*q <* 0.001, *d* = 0.54) and the out-group competition condition (*q* = 0.002, *d* = 0.43). In addition, the cooperative drumming condition showed significantly higher normalized degree than both the baseline condition (*q* = 0.042, *d* = 0.29) and the in-group competition condition (*q* = 0.042, *d* = 0.28). No other pairwise comparisons reached significance after correction (*q* s *>* 0.050). The main effect of condition was primarily driven by lower EDA connectivity in the harmonious drumming condition relative to the cooperative drumming and out-group competition conditions, together with higher connectivity in the cooperative drumming condition relative to baseline and in-group competition. This pattern partly overlapped with, but also differed from, the IBI-based results. In IBI-based networks, condition-related effects were observed for both normalized node degree and normalized node strength, with the cooperative condition showing the most pronounced increase in both the number and magnitude of cardiac synchrony-based connections. By contrast, EDA-based condition effects were restricted to normalized node degree, suggesting that EDA synchrony was primarily modulated in terms of how broadly individuals became connected to others, rather than in the overall strength of those connections. Thus, although both physiological signals were sensitive to the social task context, IBI-based networks appeared to capture changes in both the breadth and intensity of cardiac coupling, whereas EDA-based networks captured more selective changes in the breadth of electrodermal connectivity.

As observed for the IBI-based networks, experimental condition did not significantly affect the group-level organization of the EDA-based networks. No significant effects were found for network density, the normalized number of connected components, normalized clustering, degree assortativity, global efficiency, or local efficiency (*q* s *>* 0.050). Overall, condition-related differences in both physiological modalities were therefore restricted to specific node-level properties and were not accompanied by significant changes in global network topology.

## Discussion

The present study provides one of the first empirical demonstrations that physiological group synchrony can be studied as a structured topology of multiple interpersonal connections rather than as an aggregate of dyadic alignments (Carollo & Gordon, 2026). Moving beyond dyadic and aggregate approaches, we examined groups of four and five individuals engaged in a joint drumming task while cardiac and electrodermal activity were recorded simultaneously. By deriving synchrony-based networks from these signals, we show that group synchrony carries information at multiple levels of social organization: it reflects how individuals perceive the group, relates to collective behavioral coordination, varies with task context, and differs across physiological channels. This multilevel structure would be difficult to recover from approaches that average pairwise synchrony across all dyads.

The present findings can be interpreted in relation to three core features of group synchrony: flexibility, selectivity, and distribution (Carollo & Gordon, 2026). Flexibility refers to the possibility that patterns of alignment change across interaction contexts; distribution refers to the spread of coordination across multiple individuals, subgroups, and bio-behavioral systems; and selectivity refers to the possibility that individuals synchronize with particular partners, roles, or task-relevant signals while remaining decoupled from others. Flexibility was reflected in the modulation of node-level physiological connectivity across task contexts, especially during cooperation. Selectivity was reflected in the fact that synchrony was not uniformly shared across all group members, but varied in how broadly and strongly individuals were connected to others. Distribution was reflected in the organization of synchrony across multiple dyadic relations and physiological channels, as shown by the distinct patterns observed for IBI- and EDA-based networks and by the associations between EDA network topology and collective behavioral synchrony. The present study uses synchrony-based networks to illustrate how these properties can be expressed in physiological group coordination.

The social relevance of physiological group synchrony was most evident in EDA-based networks. Node-level EDA metrics, but not IBI metrics, were associated with participants’ subjective evaluations of the group, consistent with previous work linking autonomic synchrony to trust-related expectations, cooperative success, group satisfaction, and cohesion (Behrens et al., 2020; Chikersal, Tomprou, Kim, Woolley, & Dabbish, 2017; Gordon et al., 2020; Mitkidis, McGraw, Roepstorff, & Wallot, 2015; Tomashin et al., 2022). Participants who reported greater trust were both more broadly connected to others and more strongly embedded within the electrodermal synchrony network. In other words, individuals who perceived greater trust in the group were also those who showed more numerous and stronger synchrony-based physiological connections with other group members. This finding links EDA-based synchrony to the subjective experience of trust toward others and toward the group as a whole, suggesting that electrodermal coordination may capture socially meaningful variation in how individuals become physiologically connected to specific partners during group interaction. At the same time, perceived cohesion was negatively associated with EDA-based normalized strength, indicating that stronger electrodermal coupling was not uniformly associated with more positive group evaluations. This interpretation is consistent with evidence that skin-conductance synchrony may also relate to group tension and negative affect in cooperative team tasks (Mønster, Håkonsson, Eskildsen, & Wallot, 2016). This pattern is noteworthy in light of previous work showing that IBI synchrony may be associated with greater group cohesion (Gordon et al., 2020; Tomashin et al., 2022). Together, these findings suggest that IBI and EDA synchrony may capture partly distinct dimensions of group experience. Whereas cardiac synchrony may reflect a more cohesive form of interpersonal engagement, stronger electrodermal synchrony may index heightened sympathetic attunement, arousal, or interpersonal responsiveness that does not necessarily translate into perceived cohesion.

At the group level, EDA synchrony was socially informative not only through the presence of interpersonal connections, but through their distribution and organization within the network. Previous work has shown that physiological synchrony can be associated with behavioral coordination, group cohesion, and joint performance (Elkins et al., 2009; Gordon et al., 2020; Guastello, Marra, Peressini, Castro, & Gomez, 2018; Tomashin et al., 2022). However, when synchrony is studied dyadically, or when pairwise synchrony is averaged across a group, its relational topology is necessarily obscured (Carollo & Gordon, 2026; Gordon, Wallot, & Berson, 2021). The present findings extend this perspective by showing that the relation between physiological coordination and collective behavior may depend on how synchrony is organized across multiple individuals. Greater behavioral synchrony during drumming was associated with lower normalized clustering but higher local efficiency in EDA-based networks. Collective rhythmic coordination was therefore not simply related to denser physiological coupling or to the formation of tightly clustered subgroups. Rather, behavioral synchrony was associated with a local network structure in which physiological connections were efficiently organized without becoming overly segregated into closed clusters. This interpretation is consistent with hyperscanning work showing that interbrain network topology, including efficiency-based network measures, can distinguish socially coordinated interaction and team-focused coordination (Duan et al., 2015; Liu, Duan, Dai, Pelowski, & Zhu, 2021; Toppi et al., 2015). In this sense, distributed group synchrony may reflect a relational configuration in which physiological coordination balances integration and differentiation: group members remain sufficiently connected to support collective coordination, while avoiding excessive clustering that could fragment the group into isolated subgroups. Such a balance may be particularly important for collective behavior, which requires individuals to align with others while maintaining enough flexibility to adapt to the evolving dynamics of the group.

Physiological group synchrony was also flexibly organized across task contexts. This flexibility was expressed most clearly through changes in individual node embeddings rather than through detectable large-scale reconfiguration of the overall network topology. In IBI-based networks, experimental condition modulated both normalized node degree and normalized node strength. The cooperative drumming condition showed the clearest increase in both the number and magnitude of cardiac synchrony-based connections, whereas node strength was also higher during the two competition conditions relative to baseline. EDA-based networks also showed condition-related modulation, but this was expressed selectively through normalized node degree. Thus, IBI-based networks captured task-related changes in both the number and strength of cardiac connections, whereas EDA-based networks captured more selective task-related changes in the breadth of electrodermal connectivity. These condition effects extend previous work showing that physiological synchrony varies as a function of the cooperative or competitive nature of the task (Behrens et al., 2020; Chanel, Kivikangas, & Ravaja, 2012; Danyluck & Page-Gould, 2019). In dyadic approaches, such effects have necessarily been examined primarily in terms of the magnitude of the connection between two individuals. By contrast, a group-level network approach makes it possible to ask not only whether synchrony increases or decreases, but also how synchrony is selectively distributed across individuals. In this sense, the finding that cooperation modulated node-level connectivity is theoretically informative: it suggests that cooperative engagement did not merely intensify physiological coupling, but also broadened the set of partners with whom individuals became physiologically connected. The absence of significant condition effects on group-level metrics further suggests that cooperation and competition shaped how individuals were embedded within the synchrony network without substantially altering the overall architecture of the group.

The different patterns observed for IBI- and EDA-based networks may help explain why previous research on physiological synchrony and socially oriented outcomes has produced heterogeneous findings, with a large proportion of reported effects being null (Gordon & Bartsch, 2026). Physiological synchrony is often treated as a unitary construct, despite being derived from biological systems that may carry different social meanings. The present findings suggest that cardiac and electrodermal synchrony should not be considered interchangeable indices of physiological synchrony. IBI-based synchrony appeared especially sensitive to shared rhythmic engagement and cooperative coupling, whereas EDA-based synchrony was more closely related to subjective social experience and collective behavioral coordination. Thus, when the aim is to capture socially meaningful aspects of group interaction, EDA-based synchrony may provide a particularly informative physiological channel.

Together, this work supports synchrony-based network analysis as a methodological framework for characterizing group synchrony at multiple levels. By representing physiological synchrony as a coordination topology, this approach makes it possible to examine how synchrony is distributed across individuals, selectively organized across relationships, and flexibly modulated by social context. This cannot be fully captured by averaging pairwise synchrony values across all dyads, because such averages obscure who is connected to whom, how strongly, and how these connections are arranged within the group. The present findings suggest that physiological group synchrony is a multichannel and multilevel property of social coordination. Cardiac and electrodermal synchrony appear to contribute differently to this organization, with IBI-based networks capturing shared rhythmic and cooperative coupling, and EDA-based networks capturing socially meaningful variation in individual perception and collective behavior.

### Limitations

Some limitations should be considered when interpreting the results of the present study. First, the experimental conditions were administered in a fixed order. This design ensured that all groups completed the task sequence in the same way that builds on their experience in the lab, but it also means that condition effects cannot be fully separated from possible order-related influences, such as practice, fatigue, habituation, or increasing familiarity among group members. Future studies using counterbalanced or randomized task orders will be important for isolating condition-specific effects more directly.

Second, the groups examined here consisted of four or five participants. This group size was appropriate for constructing synchrony-based networks while maintaining experimental control, but it also constrains the range and stability of some network metrics. Measures such as clustering, efficiency, assortativity, and connected components may behave differently in larger groups, where more complex relational structures and subgroup configurations can emerge. Future work should therefore examine whether similar network patterns are observed in larger and more heterogeneous social groups.

Third, the synchrony networks were constructed at the condition level. This approach allowed us to compare network organization across experimental contexts, but it does not capture the moment-to-moment dynamics through which synchrony-based connections emerge, dissolve, and reorganize during interaction. This is particularly relevant given the theoretical view of group synchrony as flexible, selective, and distributed. Future studies could extend the present approach using time-varying or dynamic network analyses to examine how physiological group synchrony changes within each interaction.

Finally, the study focused on IBI and EDA as indices of physiological coordination. These signals provide complementary information about cardiac and sympathetic arousal dynamics, but they do not exhaust the range of bio-behavioral systems through which group synchrony may be expressed. Future studies could examine whether behavioral, neural, hormonal, respiratory, or additional autonomic signals show distinct network topologies, and whether these modalities interact during group coordination.

## Conclusion

The present study shows that physiological group synchrony can be studied as a structured coordination topology rather than as an aggregate of dyadic alignments (Carollo & Gordon, 2026). By preserving the configuration of physiological alignment across individuals, synchrony-based network analysis captures multilevel information about group experience that is lost when synchrony is reduced to an average of dyadic scores. This framework therefore provides a route toward studying group synchrony as a flexible, selective, and distributed property of social coordination.

The findings suggest that the social meaning of physiological synchrony depends not only on how much individuals align, but on how alignment is distributed across people, relationships, and biological systems. Cardiac and electrodermal networks contributed differently to this organization, indicating that physiological group synchrony is not a unitary phenomenon but a multichannel and multilevel property of group coordination. Synchrony-based network analysis therefore offers a methodological foundation for moving beyond whether individuals synchronize, toward understanding how patterns of synchrony are organized and what they reveal about collective social behavior.

## Author Contributions

Conceptualization: AC, IG; Data curation: AC, MS, NG, OP, EC; Formal analysis: AC, MS, NG, OP, EC, AB; Funding acquisition: IG; Investigation: MS, NG, OP, EC; Methodology: AC; Supervision: IG; Writing–original draft preparation: AC; Writing–review and editing: AC, MS, NG, OP, EC, AB, IG.

## Data availability

Data will be available upon request to the corresponding author.

## Ethics approval and consent to participate

The study was conducted in accordance with the ethical principles of the Declaration of Helsinki and was approved by the Human Subjects Institutional Review Board at Bar-Ilan University (protocol no. 221224397). All participants provided informed consent prior to participation.

